# Oral bioavailability of a noncoding RNA drug, TY1, that acts on macrophages

**DOI:** 10.1101/2024.04.27.591474

**Authors:** Shukuro Yamaguchi, Kazutaka Miyamoto, Xaviar M. Jones, Alessandra Ciullo, Ashley Morris, Kara Tsi, Eduardo Marbán, Ahmed G.E. Ibrahim

**Author notes:** Corresponding author. Ahmed G.E. Ibrahim, PhD, Smidt Heart Institute, Cedars-Sinai Medical Center, 8700 Beverly Blvd, 1090 Davis Bldg., Los Angeles, CA 90048.

## Abstract

All approved RNA therapeutics require parenteral delivery. Here we demonstrate an orally bioavailable formulation wherein synthetic noncoding (nc) RNA, packaged into lipid nanoparticles, is loaded into casein-chitosan (C2) micelles. We used the C2 formulation to deliver TY1, a 24-nucleotide synthetic ncRNA which targets the DNA damage response pathway in macrophages. C2-formulated TY1 (TY1^C2^) efficiently packages and protects TY1 against degradative enzymes. In healthy mice, oral TY1^C2^ was well-tolerated and nontoxic. Oral TY1^C2^ exhibited disease-modifying bioactivity in 2 models of tissue injury: 1) rat myocardial infarction, where a single oral dose of TY1^C2^ was cardioprotective, on par with intravenously-delivered TY1; and 2) mouse acute lung injury, where a single dose of TY1^C2^ attenuated pulmonary inflammation. Mechanistic dissection revealed that TY1^C2^ is not absorbed into the systemic circulation but is, instead, taken up by intestinal macrophages, namely those of the lamina propria and Peyer’s patches. This route of absorption may rationalize why an antisense oligonucleotide against Factor VII, which acts on hepatocytes, is not effective when administered in the C2 formulation. Thus, some (but not all) ncRNA drugs are bioavailable when delivered by mouth. Oral RNA delivery and uptake, relying on uptake via the gastrointestinal immune system, has broad-ranging therapeutic implications.

**Graphical Abstract:** Oral TY1^C2^ is taken up in the small intestine mainly by macrophages of the lamina propria and the Peyer’s patches. Macrophages that have taken up oral TY1^C2^ mediate disease-modifying bioactivity in 2 animal models of acute tissue injury. The blue + denotes the as-yet-uncertain mechanisms/mediators linking TY1 gut uptake to the drug’s systemic therapeutic benefits.

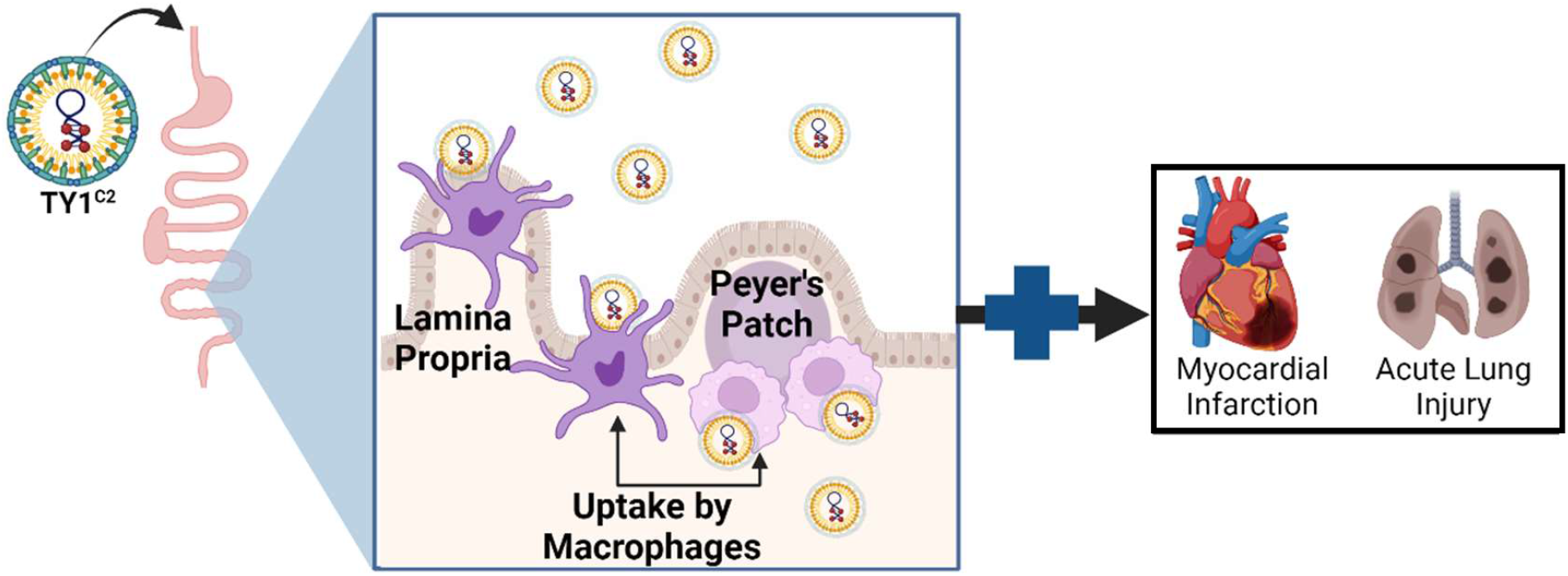

## Introduction

RNA drugs have immense potential to impact both conventional as well as novel therapeutic targets, but they face a significant hurdle: delivery. RNA drug delivery has to be effective without triggering an undesired immune reaction. Parenteral lipid nanoparticle (LNP)-packaged RNA has been met with some success, but immunogenicity remains a concern^1,2^. Systemic intravenous infusion, while less invasive than most other parenteral routes, is problematic, especially if frequent dosing is required^3^. A means to deliver RNA drugs orally, if safe and effective, is highly desirable^4^, especially for chronic disorders. However, the efficient oral delivery of fragile biotherapeutics such as RNA remains challenging, given the harsh physicochemical environment of the gastrointestinal (GI) tract: highly acidic, nuclease-rich, with an impregnable mucin barrier. We have created TY1, a 24 nucleotide chemically stabilized ncRNA drug that, when given IV, reduces tissue injury in models of myocardial infarction (MI)^5^. Here we describe a new, simple oral formulation that not only replicates the effects of intravenous TY1 on MI, but also exhibits disease-modifying bioactivity in two other disease models.

## Results

### Casein-chitosan formulation efficiently encapsulates and protects TY1

TY1 RNA, when packaged in LNP formed by DharmaFECT® (henceforth TY1-IV), is cardioprotective when administered intravenously post-MI^5^. The oral formulation consists of two additional components: casein, the primary protein in milk, and chitosan, a naturally occurring polysaccharide. Casein arose from the finding that extracellular vesicles (EVs) encapsulated in casein were orally bioactive in a mouse model of Duchenne muscular dystrophy^6^. We further added chitosan, given that the addition of a cationic polymer mediates the coagulation of casein micelles without changes in pH ^7,8^. Casein/chitosan (C2) microparticle preparations efficiently encapsulate small molecular drugs and nucleic acids^9^ in aqueous solution^7^. Biopolymers including chitosan serve to enhance drug delivery across a variety of routes including oral, nasal, intravenous, and ocular^10^. The combination of casein and chitosan forms exceptionally stable micelles for biotherapeutic delivery^11^. Leveraging these observations, we first mixed TY1-IV in a casein solution, then added medium molecular weight chitosan under acidic conditions to create TY1^C2^ (**Fig. 1A**). Nanoparticle tracking analysis of TY1^C2^ particles demonstrated a mean diameter of ∼200 nm, which was slightly larger than TY1-IV or TY1 formulated with casein only (TY1^C^; **Fig. 1B, C**), with a higher particle concentration (**Fig. 1D**). Consistent with previous reports^7^, the encapsulation efficiency of the C2 formulation was 80% (**Fig. 1E**). To assess if the C2 formulation protects against enzymatic degradation, TY1-IV, TY1^C^, or TY1^C2^ were mixed with RNase A and Proteinase K, and incubated for 20 minutes at 37°C. Only TY1^C2^ showed resistance to enzymatic degradation (**Fig. 1F**). Thus, C2 formulation efficiently encapsulates and protects TY1.

**Figure 1:**
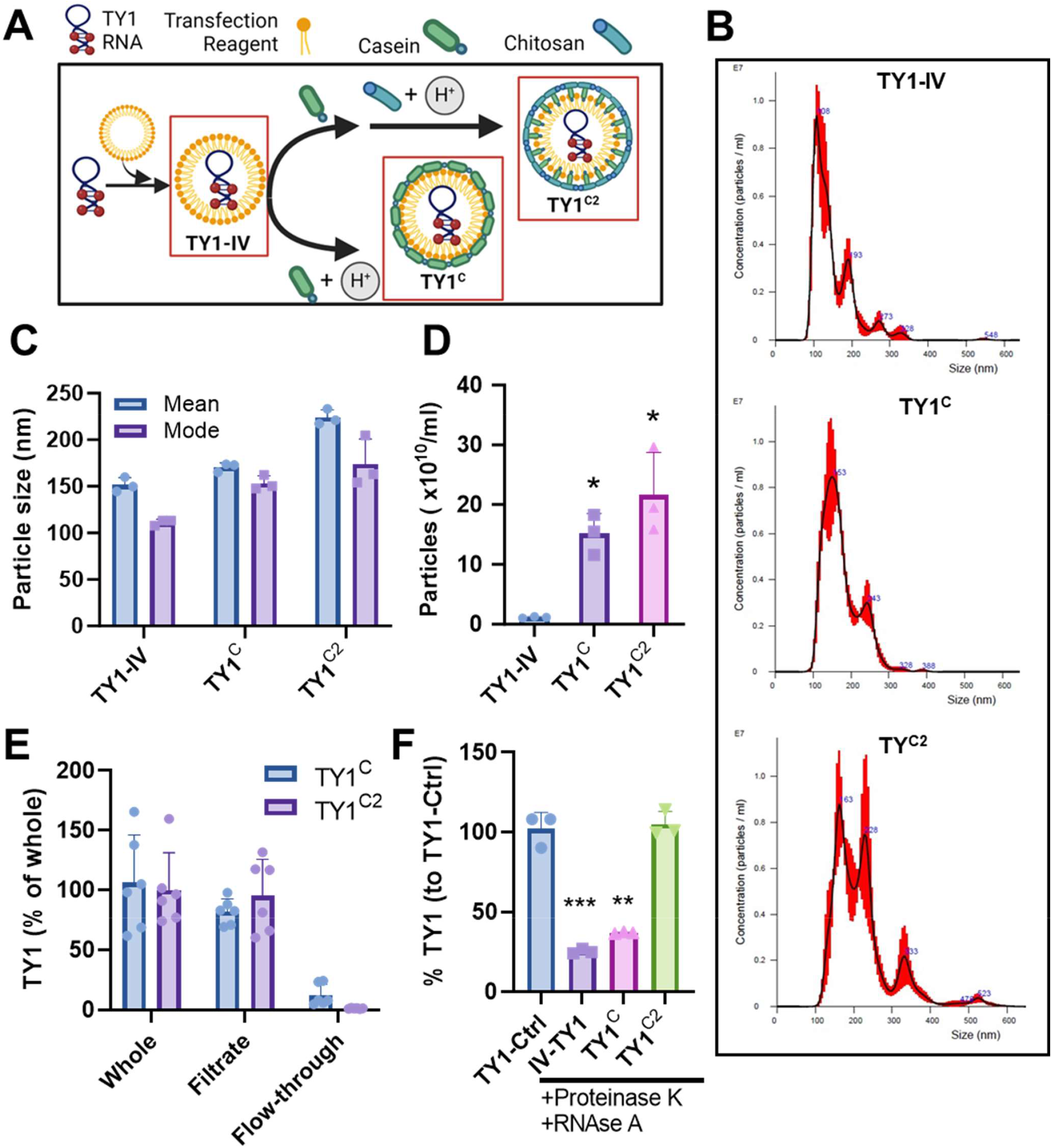
C2 Formulation of TY1. (**A**) Schematic for the formulation of TY1 in a lipid nanoparticle followed by encapsulation into casein and chitosan micelles. (**B-D**) Quantification of size and concentration of TY1^C2^ micelles by dynamic light scattering (Nanoparticle tracking analysis; n=3 biological replicates per group). (**E**) Assessment of the loading efficiency of TY1 in casein-chitosan micelles. TY1^C2^ whole formulation was filtered using a 100 kD cutoff filter to separate encapsulated (filtrate) and free (flow-through) TY1. Samples were then measured for TY1 abundance by qPCR (n= 3 biological replicates per group). (**F**) Enzyme degradation assay demonstrating resistance of TY1^C2^ to degradation by RNase A and Proteinase K (n=3 biological replicates per group). Bars represent means and error bars represent s.d. Significance was determined by one-way ANOVA; *P<0.05; **P<0.01; ***P<0.001.

### Chronic administration of TY1^C2^ is well tolerated in healthy animals

To assess toxicity and tolerability, we gavaged healthy wild-type mice with various formulations of TY1. These formulations contained various constituents of the C2 formulations to isolate any possible toxic effects of each excipient (transfection reagent, casein, or chitosan). Thus we gavaged mice twice-weekly with TY1-IV (TY1 with DharmaFECT), TY1^C^ (TY1 with DharmaFECT and casein), or the full formulation, TY1^C2^ (TY1 with DharmaFECT, casein, and chitosan)^5^. All groups were given 0.2 mg/Kg, a dose close to the previously identified IV dose of 0.15 mg/Kg^5^ but increased slightly to offset the 80% loading efficiency in C2 micelles. After 4 weeks of exposure, animals were evaluated for exercise capacity, weighed, and then euthanized for blood and tissue collection (**Fig. 2A**). At endpoint, the groups were comparable in terms of weight (**Fig. 2B**) and exercise capacity (**Fig. 2C**; the enhanced exercise capacity in TY1^C^ mice is likely spurious, given the lack of therapeutic efficacy seen below with this formulation). Furthermore, no significant changes were seen in blood chemistry, lipid panel, or liver enzymes (**Supplemental table 1**). Histological analyses of heart, lung, liver, kidney, and spleen revealed no increases in inflammatory infiltration or fibrosis (representative H&E [**Fig. 2D**] and Masson’s trichrome stained images [**Fig. 2E**], and pooled data [**Fig. 2F-I**]). Thus, none of the constituents of the TY1^C2^ formulation elicited toxic effects; chronic TY1^C2^ administration is well-tolerated in healthy animals.

**Figure 2:**
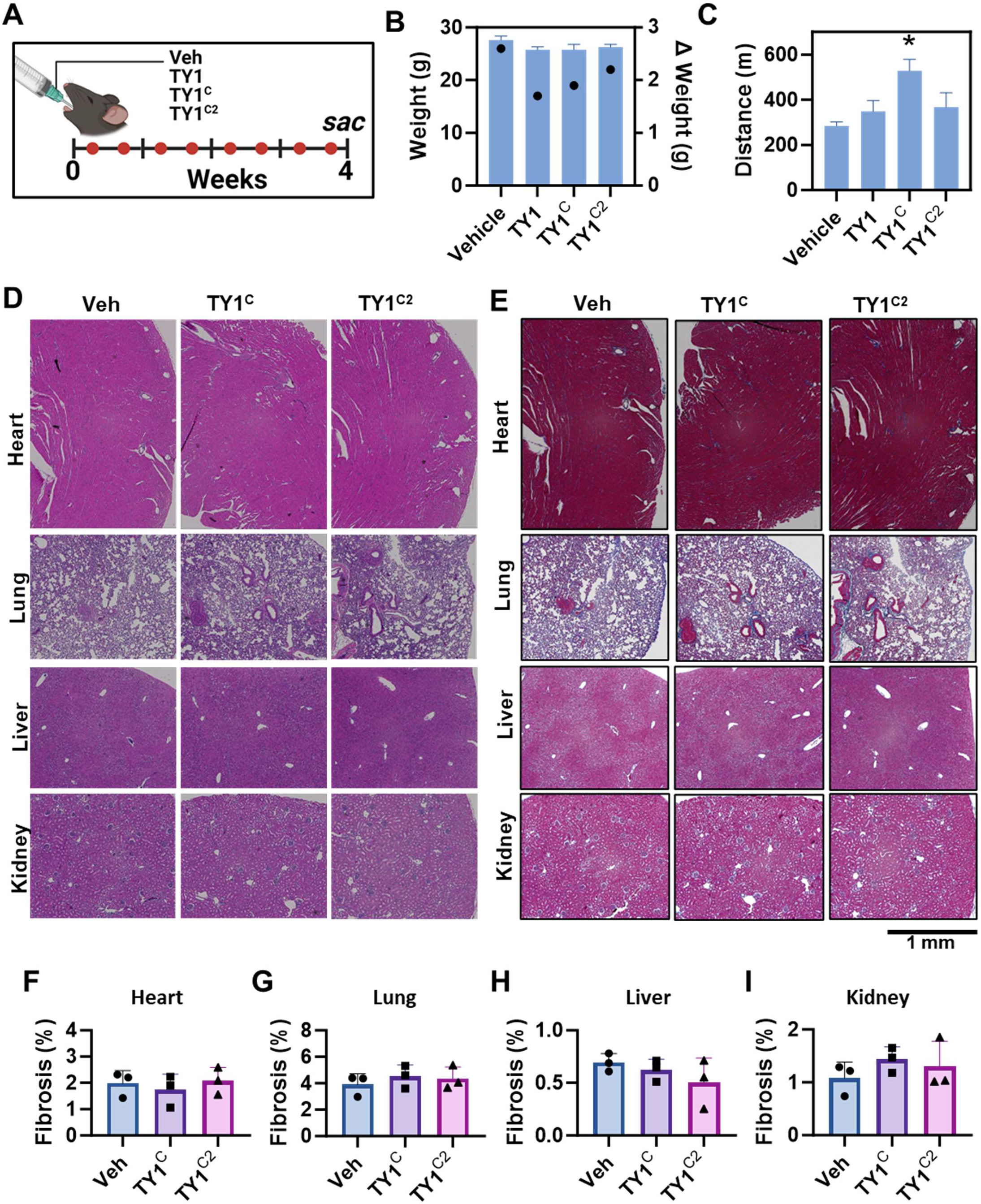
Long-term oral administration of TY1^C2^ is well tolerated in healthy animals. (**A**) Schematic for assessing the toxicity of multiple administrations of variations of TY1 formulations in healthy animals (n=5 animals per group). C57BL/6 mice were fed vehicle (PBS) or TY1^C2^ (0.2 mg/kg) twice a week for 4 weeks (eight total doses) followed by euthanasia. Other groups of animals were given TY1 formulated without chitosan (0.2 mg/kg; TY1^C^) or TY1 packaged in lipid nanoparticles (0.15 mg/Kg; TY1-IV). (**B**) Average animal weights for each group (left axis: bars) and change in body weight (left axis: dots). (**C**) Exercise endurance as measured by a treadmill test. (**D, E**) Histological analysis of animal organs four weeks post-exposure to assess inflammation (H&E; **D**) and fibrosis (Masson’s trichrome; **E**, pooled data in **F**-**I**). Bars represent means and error bars represent s.d. Significance was determined by one-way ANOVA; *P<0.05.

### TY1^C2^ protects heart and lung tissue post-injury

Intravenous TY1-IV reduces myocardial damage in multiple models of MI^5^. To compare the efficacy of oral administration, a single oral dose of TY1^C2^ or vehicle (Veh^C2^, i.e., the C2 formulation without RNA) was delivered by gavage in a rat model of MI, 20 min after reperfusion (**Fig. 3A**). Intravenous TY1-IV, also administered 20 min after reperfusion, served as a positive control^5^. Animals that received TY1^C2^ orally showed reductions of the circulating ischemic biomarker cardiac troponin I (compared to Veh^C2^; **Fig. 3B**) and infarct size (**Fig. 3C, D**). Gavage with a casein-only formulation (i.e., TY1^C^) yielded an inconsistent therapeutic benefit, unlike the robust efficacy of oral TY1^C2^, which, remarkably, is equivalent to that of intravenous TY1.

**Figure 3:**
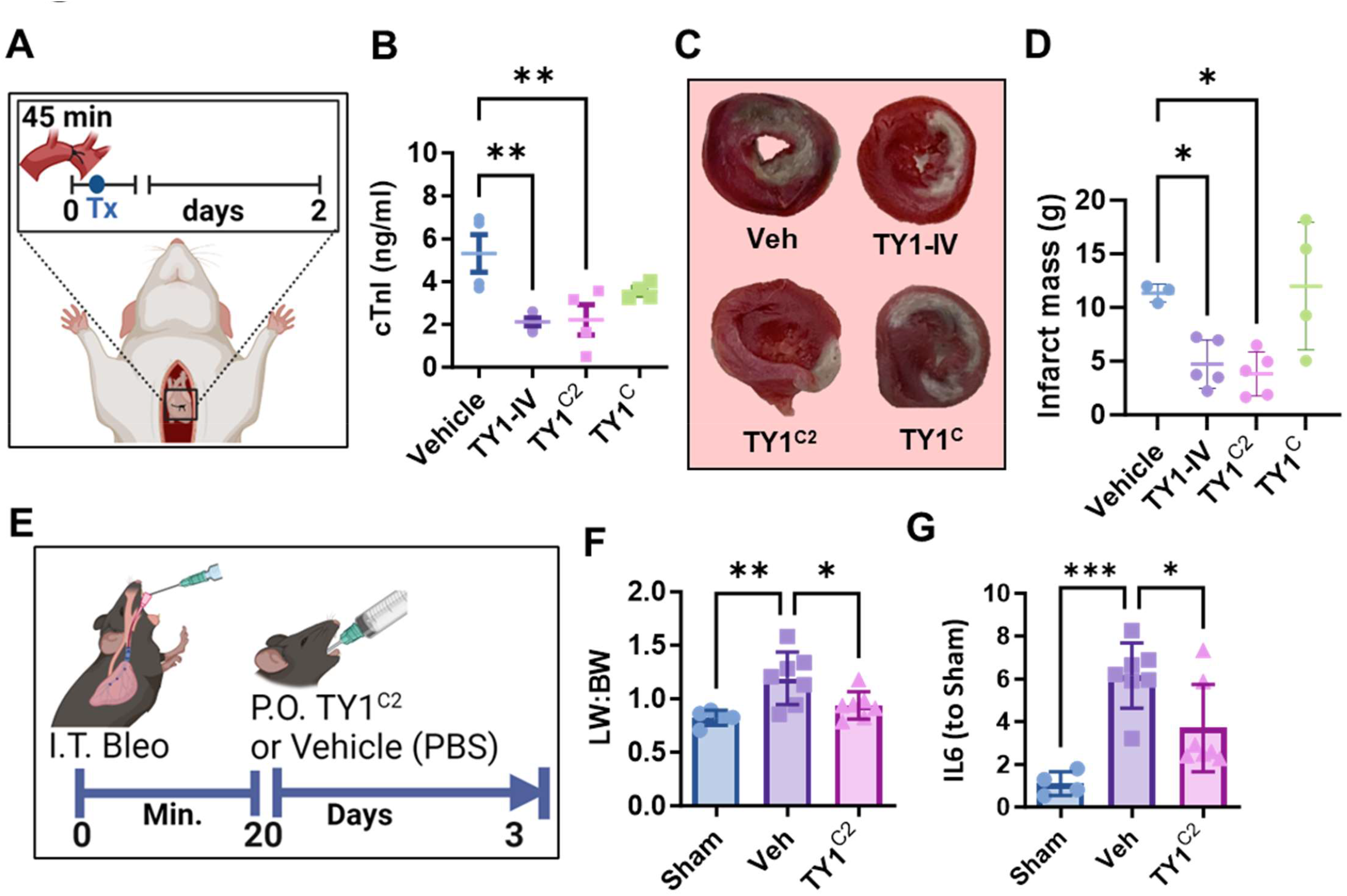
TY1^C2^ exerts disease-modifying bioactivity in models of acute organ damage. (**A**) Schematic for the rat model of myocardial infarction. Rats were subjected to ligation of the left descending coronary artery for 45 minutes. Twenty minutes after reperfusion animals received vehicle (oral), TY1-IV (intravenous), TY1^C2^ (oral), or TY1 formulated with LNPs and casein only (oral; TY1^C^, n=4-5 rats per group). Forty-eight hours post-injury, animals fed TY1^C2^ had reduced cardiac tissue damage as demonstrated by reduced circulating levels of cardiac troponin as measured by ELISA (**B**) and infarct size as quantified by TTC staining (representative images; **C** and pooled data; **D**). (**E**) Schematic for the mouse model of bleomycin-induced acute lung injury (n=5-7 mice per group). 10-week-old C57BL/6 mice received a single intratracheal administration of saline (sham) or bleomycin sulfate (0.7 mg/kg). Twenty minutes later, injured animals received oral vehicle (casein-chitosan micelles only), or TY1^C2^ (0.2 mg/kg). Seventy-two hours later, animals that had received a single oral dose of TY1^C2^ showed reduced lung inflammation as measured by lung weight to body weight ratio (**F**) and IL6 expression in lung tissue as measured by qPCR (**G**). Bars represent means and error bars represent s.d. Significance was determined by one-way ANOVA; *P<0.05; **P<0.01; ***P<0.001.

To determine if TY1^C2^ can exert broad disease-modifying bioactivity, we investigated TY1^C2^ in a bleomycin-induced acute lung injury model. Seven-to 10-week-old mice received a single intratracheal instillation of bleomycin sulfate followed by a single oral gavage of vehicle or TY1^C2^ twenty minutes later (**Fig. 3E**). Lung tissue isolated from TY1^C2^ animals showed attenuated inflammation as measured by reduced lung weight to body weight ratio (**Fig. 3F**) and attenuated expression of Il6, a pro-inflammatory cytokine (**Fig. 3G**).

### Orally-delivered TY1^C2^ is taken up by macrophages in the small intestine

A qPCR-based method is useful for tracking the biodistribution of EVs by amplifying NT4, a naturally occurring RNA with a sequence 96% identical to that of TY1^12^. We used the same method here to track the biodistribution of orally-delivered TY1^C2^. Despite the sensitivity of this technique^5^, TY1 was neither detectable in tissue nor in plasma (**Supplementary Fig. 1A**). TY1 remained undetected even when the administered dose was increased 100-fold, raising the possibility that orally administered TY1^C2^ is not absorbed into the circulation (**Supplementary Fig. 1B**). To investigate whether another orally-administered RNA can enter the systemic circulation, we used an siRNA against coagulation factor VII (siFVII), which is produced by hepatocytes, the cell target for intravenously or intraperitoneally-injected RNA^12^. Twenty-four hours after animals were injected intravenously or intraperitoneally with LNP-packaged siFVII, liver tissue exhibited potent dose-dependent suppression of Factor VII (**Supplementary Fig. 1C, D**). However, liver tissue from animals that had received oral siFVII formulated in C2 showed no suppression of Factor VII, even at doses much higher than those effective parenterally (**Supplementary Fig. 1C, E**). Similarly, an siRNA against a more ubiquitous gene target, Gapdh (siGap), worked well parenterally but not orally (**Supplementary Fig. 1F, G**). Thus, when given orally, two other small RNAs formulated in C2 had no systemic effects. All three compounds tested—TY1, siFVII, and siGap—are undetectable in plasma when given orally in the C2 formulation, but oral TY1 is unique among these in exerting strong physiological effects *in vivo*, comparable to those of intravenous delivery. Thus, the C2 formulation does not provide oral access of its payload to circulating plasma, but something about TY1^C2^ nevertheless renders it orally bioavailable.

While siFVII and siGap block translation of their targeted genes in hepatocytes, TY1 is active in a different cell type: macrophages. TY1 is taken up by macrophages, which are necessary and sufficient to confer the therapeutic benefits of TY1^5^. We therefore investigated the possibility that oral TY1^C2^ might be taken up by macrophages in the GI immune system, specifically those in the small intestine. Animals fed TY1^C2^ had unique enrichment of TY1 (as measured by qPCR) in Peyer’s patches and, to a lesser extent, in nearby intestinal tissue (**Supplementary Fig. 2A**). Another, distant secondary lymphoid organ, the spleen, showed no TY1 (**Supplementary Fig. 2A**). This opened the possibility for visualizing fluorescent-labeled TY1^C2^. A pilot study evaluating signal intensity at one hour post-feeding (as informed by qPCR data in **Supplementary Fig. 2A**) demonstrated the feasibility of this approach (**Supplementary Fig. 2B**). Tracking fluorescent Alexa Fluor 750-tagged TY1^C2^ (^A750^TY1^C2^), small intestine isolated from animals fed ^A750^TY1^C2^ showed a strong signal at 30 minutes, which waned by four hours (**Fig. 4A, B**). Gut-associated lymphoid structures, including Peyer’s patches, are inductive sites where intestinal contents interface with the immune system^13^. Among cells isolated from the small intestine one hour post-gavage, intestinal epithelial cells showed no TY1 uptake (**EpCAM; Supplementary Fig. 3A, Fig. 4C, D**). In contrast, macrophages (F4/80^+^) exhibited robust TY1 signal in both Peyer’s patches (**Supplementary Fig. 3B, Fig. 4E, F**) and in the lamina propria (**Supplementary Fig. 3C, Fig. 4G, H**). Thus, orally administered TY1^C2^ is taken up by macrophages in the small intestine. Somehow, by as-yet-undetermined mechanisms, this localized GI lymphoid uptake leads to reparative benefits in distant tissue, as seen in myocardial infarction and acute lung injury (**Fig. 3**).

**Figure 4:**
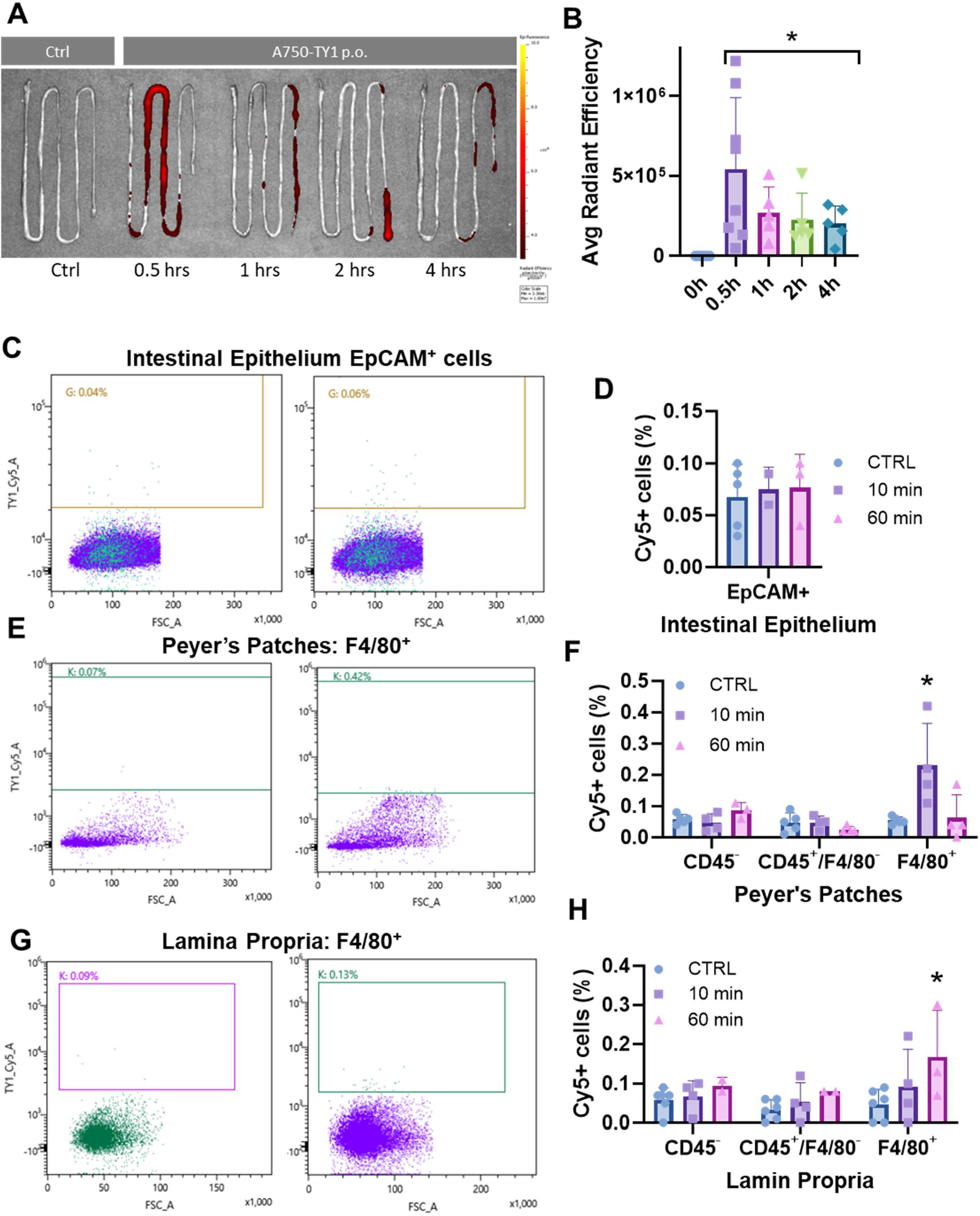
Intestinal macrophages take up TY1^C2^ after oral delivery. (**A**) *Ex vivo* imaging of small intestine from mice fed TY1^C2^ made with Alexa Fluor 750-tagged TY1 (^A750^TY1^C2^) at 30-, 60-, and 120-minutes post-gavage. (**B**) Pooled data of TY1C2 signal (expressed as average radiant efficiency) in the small intestine over time (n=5-8 mice per group). Flow cytometry analysis of ^A750^TY1^C2^ uptake by intestinal epithelial cells (EpCAM^+^; **C, D**), Peyer’s patch cells (**E, F**), and lamina propria cells (**G, H**; n=4 animals per group). Bars represent means and error bars represent s.d. Significance was determined by one-way ANOVA; *P<0.05.

## Discussion

Oral delivery of RNA therapeutics is fraught with challenges. RNA molecules and the lipid nanoparticles that encapsulate them can be degraded by acid in the stomach and nucleases in the gastrointestinal tract. Furthermore, RNA molecules, owing to their size and charge, are not able to permeate the intestinal epithelium, limiting bioavailability^14^. Efforts to overcome these challenges have focused on chemical modifications of RNA and nanoparticle carriers to enhance their stability. Improvements in RNA include ribose modifications and cholesterol conjugates, while various nanoparticle excipients have been explored, with limited success^15^. Regardless of chemical modifications, unprotected RNA molecules are largely degraded as they pass through the GI tract. Moreover, even if nanoparticles provide sufficient protection of RNA molecules, mucous penetration, and subsequent absorption through the tight gap junctions of intestinal villi remain as barriers^16^.

Here we demonstrate a polymer-based formulation of a small non-coding RNA with disease-modifying bioactivity, TY1, for oral delivery. We found the C2 formulation to efficiently encapsulate TY1 and protect the oligonucleotide from the harsh GI environment. TY1^C2^ is well-tolerated in healthy animals after chronic exposure. In two models of acute organ injury, a single oral administration of TY1^C2^ demonstrated tissue-protective activity equivalent to that of intravenous TY1. Surprisingly, C2 formulation does not deliver free RNA cargo to the systemic circulation, as gauged by plasma concentration. Instead, RNA cargo is taken up and sequestered by macrophages in the lamina propria and Peyer’s patches.

Given the importance of macrophages—they are both necessary and sufficient for TY1 efficacy^5^— it stands to reason that the intestinal macrophages that take up TY1^C2^ initiate the systemic therapeutic effect. However, the mechanism by which this happens remains to be elucidated. A recent study of another orally delivered RNA proposed a relay system involving, in sequence, intestinal macrophages in the Peyer’s patches, lymphocytes, dendritic cells, and systemic circulation^17^, but direct experimental evidence was lacking. Other studies suggest that macrophages in the intestine, once activated by immunostimulatory nanoparticles like yeast-derived glucans, stimulate uptake by macrophages^18^. Efficient oral delivery of siRNA targeting Map4K4 to intestinal macrophages has been reported to suppress systemic inflammation in a model of LPS-induced sepsis; macrophages with suppressed Map4k4 were detected in distant tissues including lung, spleen, muscle, liver, and the peritoneum^19^. Similarly, chitosan has been shown to exert immunostimulatory properties^20,21^ and systemically deliver orally-administered-siRNA^22^. However, challenges with using chitosan exclusively for oral formulation include limited permeability across the intestinal epithelium^23^. Conversely, casein facilitates uptake by cells and enhances sustained bioavailability^24^. Thus, when combined, the immunostimulatory properties of chitosan and bioavailability-enhancing properties underlie the C2 formulation’s exceptional ability to deliver small RNA cargo to intestinal macrophages with effects that are, dose-for-dose, comparable to those of IV administration. The C2 formulation may, more broadly, enable the targeting of macrophages as a therapeutic strategy, not only for TY1 but also for other drugs that act on macrophages.

## Methods

### Nanoparticle Tracking Analysis

The size and concentration of TY1^C2^ nanoparticles were characterized using nanoparticle tracking analysis by NanoSight (Malvern Panalytical, Malvern, UK).

### Experimental Animals

All studies were performed at Cedars-Sinai Medical Center in accordance with the Institutional Animal Care and Use Committee guidelines. Six to ten-week-old male or female C57BL/6J mice (Jackson Laboratory, Maine, USA) were used for safety evaluation, biodistribution study, and bleomycin-induced lung injury model. Seven to 10-week-old Wistar-Kyoto female rats (Envigo, Indiana, USA) were used for the MI model.

### TY1 Intravenous (TY1-IV) injection

Animals were treated with TY1 oligo ribonucleotide (0.15 mg/kg body weight) after adding DharmaFECT1® (0.36 ml/kg body weight, T-2001, Horizon Discovery) by retro-orbital injection. Injections were performed on alternating eyes (no more than 3 injections per eye) under general anesthesia; no signs of ocular injury were observed.

### TY1 oral formulation and gavage

Each animal was fed TY1-IV (TY1 in lipid nanoparticles [Dharmafect®]) formulated in casein-chitosan (C2). Briefly, TY1-IV (0.2 mg/kg body weight) was mixed with a 5% casein solution from bovine milk (125 μl per mouse, C4765, Millipore Sigma) and incubated for 15 minutes at room temperature with gentle agitation. Following incubation, 100μl of a 0.1% acetic acid solution (695092, Sigma-Aldrich) / 0.2% chitosan (448869, Millipore Sigma) was added dropwise and mixed well. After 60 min incubation at room temperature, the solution was administered via an oral gavage needle (01-290-3B, Fisher Scientific); the animal was held with the body tilted upward, and the needle was inserted into the mouth gently.

### Enzyme degradation assay

TY1-IV, TY1^C^, and TY1^C2^ were mixed with 5 μg/ml of RNase A (19101, QIAGEN) and 1 mg/ml of Proteinase K (EO0491, Fisher Scientific), and incubated for 20 minutes at 37°C using a tube rotator to gently mix samples during incubation. As a control, TY1-IV without RNase A and Proteinase K was incubated in the same manner. TY1 was purified from samples using miRNeasy Mini Kit (217004, QIAGEN) according to the manufacturer’s protocol with minor modifications. Briefly, samples were lysed and homogenized by adding 700 μl of QIAzol and vortexing for 30 seconds. Spike-In Control (5.6 x 10^8^ copies, 219610, QIAGEN) was added to the samples for assessment of recovery of small RNAs during the process. Samples were then mixed with 140 μl of chloroform and centrifuged at 12,000 x g for 15 minutes at 4°C. After centrifugation, the upper aqueous phase was mixed with 1.5 volumes of 100% ethanol and transferred to an RNeasy Mini spin column. Washing steps using Buffer RWT/RPE were performed according to the manufacturer’s protocol. 40 μl of RNase-free water was added to the column membrane and the column was centrifuged at 8,000 x g for 1 minute to elute the RNA. RT-qPCR was performed as described below.

### Toxicity study

Safety evaluation was conducted by administrating TY1^C2^ to healthy mice orally twice a week for 4 weeks. Exercise endurance (details below) and body weight were measured at the endpoint followed by isolation of organ samples. Harvested tissues were embedded in paraffin and cut into sections, which were stained with hematoxylin and eosin or Masson’s trichrome. Blood chemistry tests were performed by Antech (California, USA).

### Rat Model of Myocardial Infarction

All rats were housed in a pathogen-free facility (cage bedding: Sani-Chips, PJ Murphy) with a 14 hours/10 hours light/dark cycle with food (PicoLab Rodent Diet 20 [no. 5053], LabDiet) and water provided ad libitum. *In vivo*, experimental protocols were performed on 7 to 10-week-old female Wistar-Kyoto rats. To induce MI, a thoracotomy was performed at the fourth intercostal space to expose the heart under general anesthesia. A 7–0 silk suture was then used to ligate the left anterior descending coronary artery, which was removed after 45 min to allow for reperfusion. Twenty minutes later, animals received vehicle (PBS only) or TY1^C2^ (0.2 mg/Kg as described previously) or TY1 formulated only in casein (TY1^C^; 0.2 mg/Kg). As a positive control, a group of animals received TY1-IV (0.15 mg/Kg) injected in the retro-bulbar space.

### TTC staining

Two days post-MI, 10% KCl was injected into the left ventricle to arrest hearts in diastole. Then, hearts were harvested, washed in PBS, and cut into 1‐mm sections from apex to base, above the infarct zone. Sections were incubated with 1% 2,3,5‐triphenyl‐2H‐tetrazolium chloride solution (TTC, Sigma-Aldrich) for 30 minutes at 37°C in the dark and washed with PBS. Then, sections were imaged and weighed. The infarcted zones (white) were delineated from viable tissue (red) and analyzed (ImageJ software). Infarct mass was calculated in the tissue sections according to the following formula: (infarct area/total area) / weight (mg).

### ELISA

Blood was collected from animals at the study endpoint in EDTA tubes. After being left undisturbed at 4°C for 30 minutes, plasma was obtained after 15-minute centrifugation at 4,000 rpm. Cardiac TnI was quantified using the RAT cardiac troponin-I ELISA kit (cTNI-2-HSP, Life Diagnostics) according to the manufacturer’s protocol. IL-6 and BNP plasma levels were analyzed using the following ELISA kit: Mouse IL-6 Quantikine ELISA Kit (M6000B, R&D systems), Mouse BNP EIA (EIAM-BNP-1, RayBiotech) according to manufacturers’ instructions.

### Mouse model of bleomycin-induced lung injury

10-week-old female C57BL/6 mice were given a single intratracheal instillation of a 50 μl solution of bleomycin sulfate (0.7 mg/kg) under general anesthesia. Twenty minutes later, animals were orally administered 230 μl of vehicle (casein-chitosan-LNP micelles formulated without TY1) or TY1^C2^. At seventy-two hours post-injury, animals were weighed and sacrificed, and lungs were harvested for weighing and RNA isolation for subsequent qPCR to measure Il6 expression.

### Exercise endurance (treadmill) test

Mice were placed on an Exer 3/6 rodent treadmill (Columbus Instruments, Ohio, USA) at a 5-degree elevation. At first, the speed was increased by 5 m/min for 2 min, and the speed remained 10m/min for 5 min. Then the speed was increased by 2 m/ 2min until mice were exhausted, as defined as the inability of the mouse to return to the treadmill from the shock grid for 10 seconds.

### RNA isolation and RT-qPCR

Total RNA was extracted from mouse tissue using RNeasy plus kit (74136, QIAGEN) and Maxtract High-density tubes (129056, QIAGEN). cDNA was synthesized from RNA using a High-capacity cDNA reverse transcription kit (4368813, Applied Biosystems) according to the manufacturer’s protocol. Real-time PCR (QuantStudio 12K Flex Real-Time PCR system; Thermo Fisher Scientific) was performed in triplicate using the following TaqMan Gene Expression Assay probes: IL-6 (Mm00446190_m1), analyzed by the ddCt method. TY1 was extracted from mouse tissue using miRNeasy mini kit (217004, QIAGEN) or miRNeasy Serum/Plasma kit (217184, QIAGEN) according to the manufacturer’s protocol. cDNA was synthesized from RNA using miRCURY LNA RT Kit (339340, QIAGEN) according to the manufacturer’s protocol. Real-time PCR was performed in duplicate or triplicate using the following kits and primers; miRCURY SYBR Green PCR Kit (339345, QIAGEN), miRCURY LNA miRNA PCR Assay (339306, QIAGEN), miRCURY LNA miRNA Custom PCR Assay (339317, QIAGEN).

### Oral-siRNA bioavailability study

Ambion *In Vivo* siRNA targeting FVII (4459408, Fisher Scientific), Gapdh (4459407, Thermo Fisher Scientific), or vehicles were injected intravenously or intraperitoneally (0.1 to 2 mg/kg body weight) according to the manufacturer’s protocol. For oral administration, siRNAs were mixed with transfection reagent (Invivofectamine 3.0, IVF3001, Thermo Fisher Scientific) in a 1.2:1 (μg:μL) ratio followed by incubation for 30 minutes at 50°C. C2-formulated siRNAs were then made by adding 5% casein solution and 0.2% chitosan/1% acetic acid solution as previously described. Organs including the liver were harvested 24 or 72 hours after treatment with siRNAs. Livers were digested using Bead Mill 4 (Fisher Scientific), and RNA purification and cDNA synthesis were performed as described above. qPCR was performed using the following Taqman Gene Expression Assay (Thermo Fisher Scientific): Coagulation factor VII (F7) (Mm00487329_m1) and Glyceraldehyde 3-phosphate dehydrogenase (Gapdh) (Mm99999915_g1).

### In Vivo Imaging System (IVIS)

Small intestinal tissues were harvested from mice 30 minutes, 1 hour, 2 hours, and 4 hours after treatment with C2-formulated TY1 conjugated with Alexa Fluor 750. The lumen of the intestine was thoroughly flushed with PBS using a 10 ml syringe and 14-gauge catheter to get rid of wastes and any residuals inside. Intestinal tissues were then cut through longitudinally and any visible residuals at the inner lining were further removed. Tissues were then rigorously rinsed with PBS four times and stored on ice for optical imaging. A radiant efficiency ((photons/sec/cm^2^/str)/( μW/cm^2^)) of the fluorescence from the tissues was measured with an IVIS Lumina XR (Revvity, Massachusetts, USA) using 745/800 nm emission filter set with a fixed exposure time. Five to 8 samples were collected per group for quantitative analysis.

### Flow Cytometry

Small intestines were harvested from mice after treatment with C2-formulated TY1 conjugated with Cy5 or vehicle and rinsed with PBS as described above. Cells were extracted from intestinal tissues using the Lamina Propria Dissociation Kit (130-097-410, Miltenyi) to make a single-cell suspension according to the manufacturer’s protocol with minor modifications. Briefly, tissues were cut laterally into pieces of ∼0.5 cm in length and incubated with predigestion solution for 20 minutes at 37°C. Tissues were then vortexed and applied onto a 100 μm cell strainer. Flow through was kept on ice for the collection of epithelial cells. Tissues were further incubated with HBSS for another 20 minutes and flow-through after filtration with a cell strainer was kept on ice. Tissues were then incubated with enzyme mix in digestion solution for 30 minutes at 37°C. After rigorous vortexing and pipetting, the cell suspension was applied onto a 100 μm cell strainer, and flow through was centrifuged at 300 x g for 5 minutes at room temperature. The cells were resuspended in 1 ml of ACK Lysing Buffer and incubated for 1 minute at room temperature to remove erythrocytes. Cells were then pelleted and resuspended with PBS. Two x 10^6^ cells were isolated for staining. Cells were incubated with 1 μl of Ghost Dye Red 710 (13-0871, Cytek Biosciences) for 30 minutes at 4°C. Cells were pelleted and resuspended in 100 μl of Cell Staining Buffer (420201, BioLegend) followed by incubation with 1 μl of TruStain FcX™ PLUS (156603, BioLegend) for 15 minutes at 4°C to block non-specific Fc-mediated interactions. Cell suspensions were mixed with antibodies (1 μl of FITC-CD45 (11-0541-82, Thermo Fisher Scientific), 1.25 μl of PE-F4/80 (12-4801-82, Thermo Fisher Scientific), 5 μl of Alexa 594-EpCAM (BS-4889R-A594, Thermo Fisher Scientific), and 10 μl of APC-Microfold (130-102-149, Miltenyi Biotec)) and incubated for 30 minutes at 4°C. Cells were washed 2 times by resuspension in 2 ml of Cell Staining buffer and centrifugation at 800 x g for 5 minutes. Finally, cells were resuspended in 500 μl of FluoroFix Buffer (422101, BioLegend). Unstained and isotype control samples were also prepared.

### Statistical analysis

Statistical parameters including the number of samples (n), descriptive statistics (mean and standard deviation), and significance are reported in the figures and figure legends. In general, at least n = 3 was used for each time point and each experiment. Differences between groups were examined for statistical significance using the Student’s t-tests or analysis of variance. Differences with p values < 0.05 were regarded as significant.

## Author contributions

Conceptualization: AGI, EM

Methodology: SY, KY, AAM, AGI, EM

Investigation: SY, KY, XV, AAM, AC, and KT

Data analysis: SY, KY, XV, AC, AAM, and AGI

Funding Acquisition: AGI, EM

Supervision: AGI, EM

Writing: SY, AGI, EM

