## Supplementary Data for "Oral bioavailability of a noncoding RNA drug, TY1, that acts on macrophages"

#### Supplementary Figures

#### Supplementary Figure Legends

**Supplementary Table 1: Blood chemistries from animals dedicated to TY1 toxicity study.** Metabolic panel of plasma samples from healthy animals that had been given vehicle, oral TY1 packaged in LNP only, $TY1^{C}$, or $TY1^{C2}$ twice a week for four weeks (n= 5 animals per group).

**Supplementary Figure 1: C2 formulation does not deliver small RNA cargo systemically.** (**A**) qPCR demonstrated a lack of TY1 in organ tissues at 0, 20, 60, and 100 minutes post oral administration of $TY1^{C2}$ (0.2 mg/kg). (**B**) TY1 was still undetectable even when the oral dose was increased 100-fold (20 mg/kg). (**C**) Schematic for delivering intravenous and orally formulated siRNA to assess effects on liver tissue. (**D**) Successful suppression of Factor VII in liver tissue following intravenous or intraperitoneal administration of siRNA against Factor VII (siFVII). (**E**) Oral administration of C2-formulated siFVII at higher doses of siFVII failed to suppress Factor VII in the liver. (**F**) Intravenous administration of siRNA against Gapdh (siGap) led to successful suppression of Gapdh in liver tissue. (**G**) Oral administration of C2-formulated siGap failed to suppress Gapdh in both liver tissue and resident liver macrophages. Bars represent group means and error bars represent s.d.

**Supplementary Figure 2: $TY1^{C2}$ biodistribution in mouse tissue.**
(**A**) qPCR of TY1 in Peyer's patches, intestinal tissue, and spleen demonstrating absorption of $TY1^{C2}$ by Peyer's patches and (to a lesser extent) intestinal tissue (n=4 animals per group). (**B**) Detectable fluorescence signal of ${}^{A750}TY1^{C2}$ in the mouse small intestine one hour post oral delivery with notable absence in other organs.

**Supplementary Figure 3: $TY1^{C2}$ uptake by intestinal macrophages (A)** Gating strategy for assessing uptake of ${}^{A750}TY1^{C2}$ in intestinal epithelial cells (**B**), lamina propria (**C**), and Peyer's patches (**D**).

### Supplementary Table 1

| | CTRL | TY1-IV | TY1$^{C}$-Oral | TY1$^{C2}$-Oral | p-value |
|---|---|---|---|---|---|
| TP (g/dL) | 3.00 ± 0.1 | 2.86 ± 0.1 | 2.84 ± 0.1 | 2.90 ± 0.1 | 0.215 |
| Alb (g/dL) | 1.94 ± 0.1 | 1.86 ± 0.1 | 1.86 ± 0.1 | 1.92 ± 0 | 0.0568 |
| Globulin (g/dL) | 1.06 ± 0.1 | 1.00 ± 0.1 | 0.98 ± 0.1 | 0.98 ± 0.1 | 0.525 |
| AST (IU/L) | 20.6 ± 1.3 | 23.2 ± 5.7 | 20.0 ± 1.4 | 18.4 ± 1.1 | 0.139 |
| ALT (IU/L) | 13.2 ± 1.3 | 12.6 ± 1.9 | 12.2 ± 0.8 | 12.0 ± 1.0 | 0.523 |
| T-bil (mg/dL) | 0.1 ± 0 | 0.1 ± 0 | 0.1 ± 0 | 0.1 ± 0 | - |
| Cre (mg/dL) | 0.2 ± 0 | 0.2 ± 0 | 0.2 ± 0 | 0.2 ± 0 | - |
| Glucose (mg/dL) | 212.6 ± 20.8 | 208.8 ± 21.4 | 223.8 ± 20.3 | 231.4 ± 8.8 | 0.240 |
| T-Cho (mg/dL) | 60.4 ± 8.3 | 57.6 ± 6.7 | 56.8 ± 6.2 | 47.0 ± 24.7 | 0.466 |
| TG (mg/dL) | 101.4 ± 26.2 | 77.4 ± 20.9 | 76.4 ± 10.4 | 84.8 ± 30.5 | 0.329 |
| AMY (IU/L) | 409.0 ± 40.8 | 345.2 ± 37.6 | 346.0 ± 50.9 | 355.4 ± 18.5 | 0.0572 |
| CPK (IU/L) | 28.4 ± 19.7 | 38.4 ± 10.8 | 91.0 ± 98.6 | 60.6 ± 40.8 | 0.310 |
| WBC (/µL) | 4,117 ± 2,571 | 3,308 ± 1,423 | 4,317 ± 3,483 | 3,620 ± 858 | 0.915 |

### Supplementary Figure 1

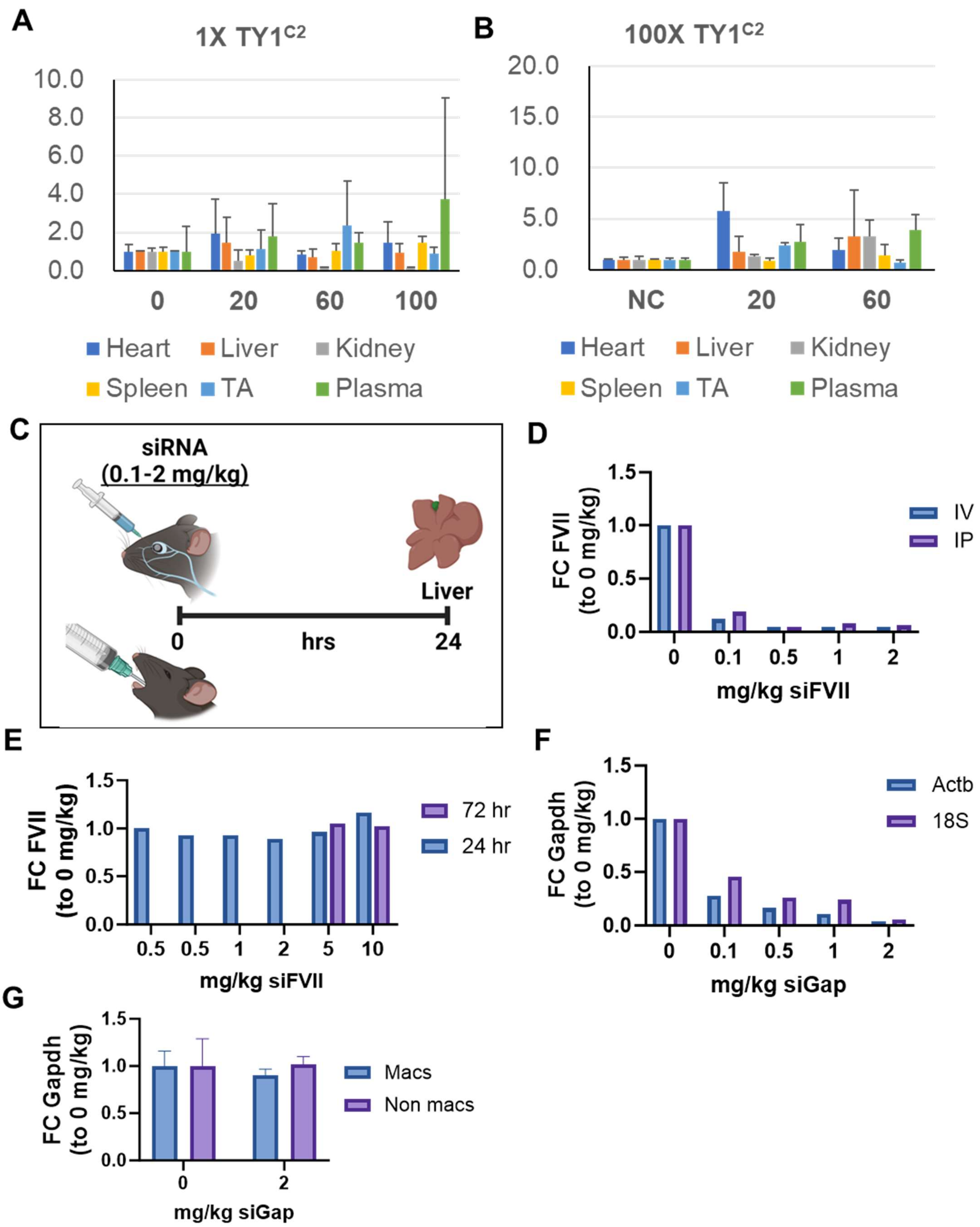

#### Supplementary Figure 2

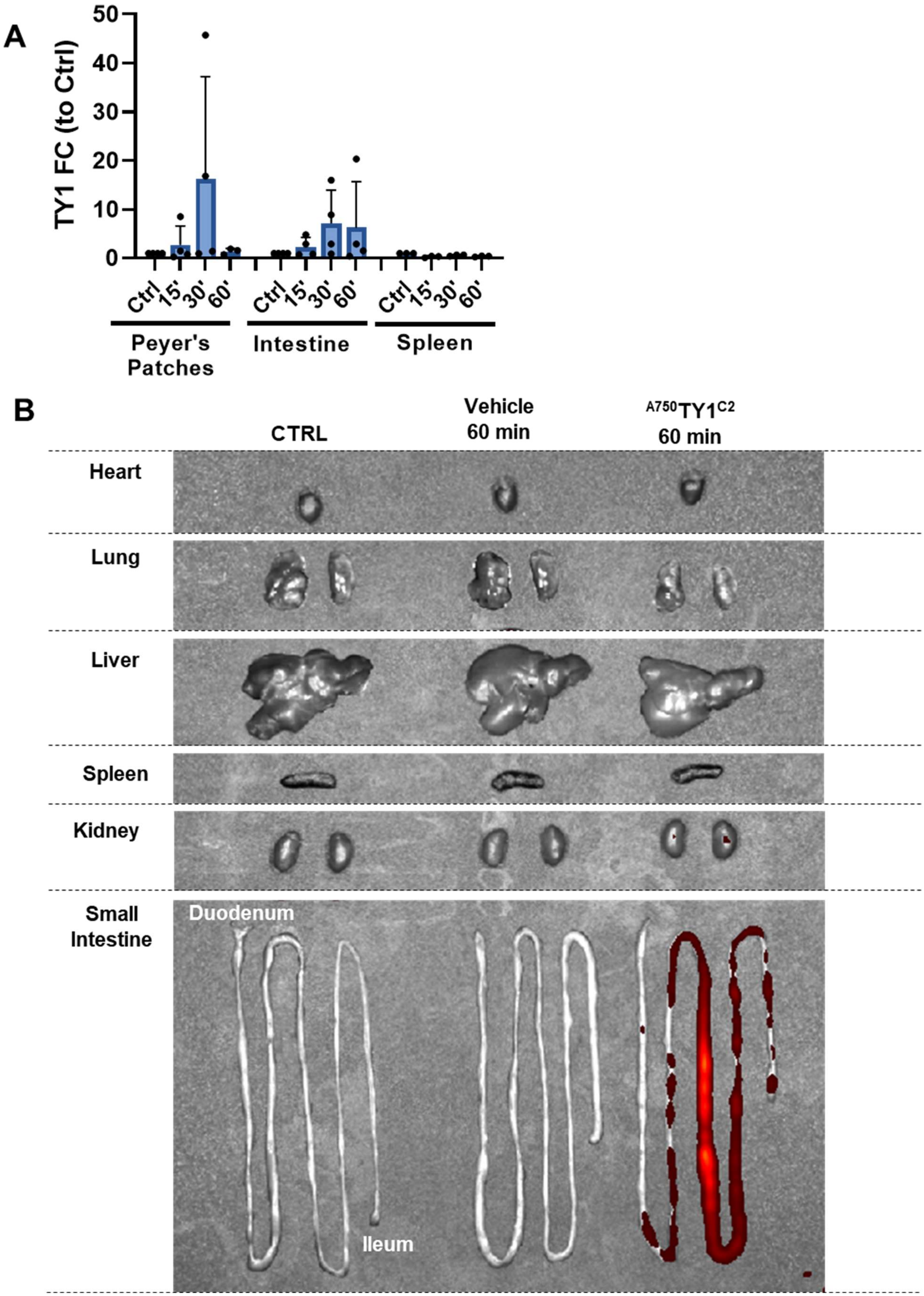

### Supplementary Figure 3

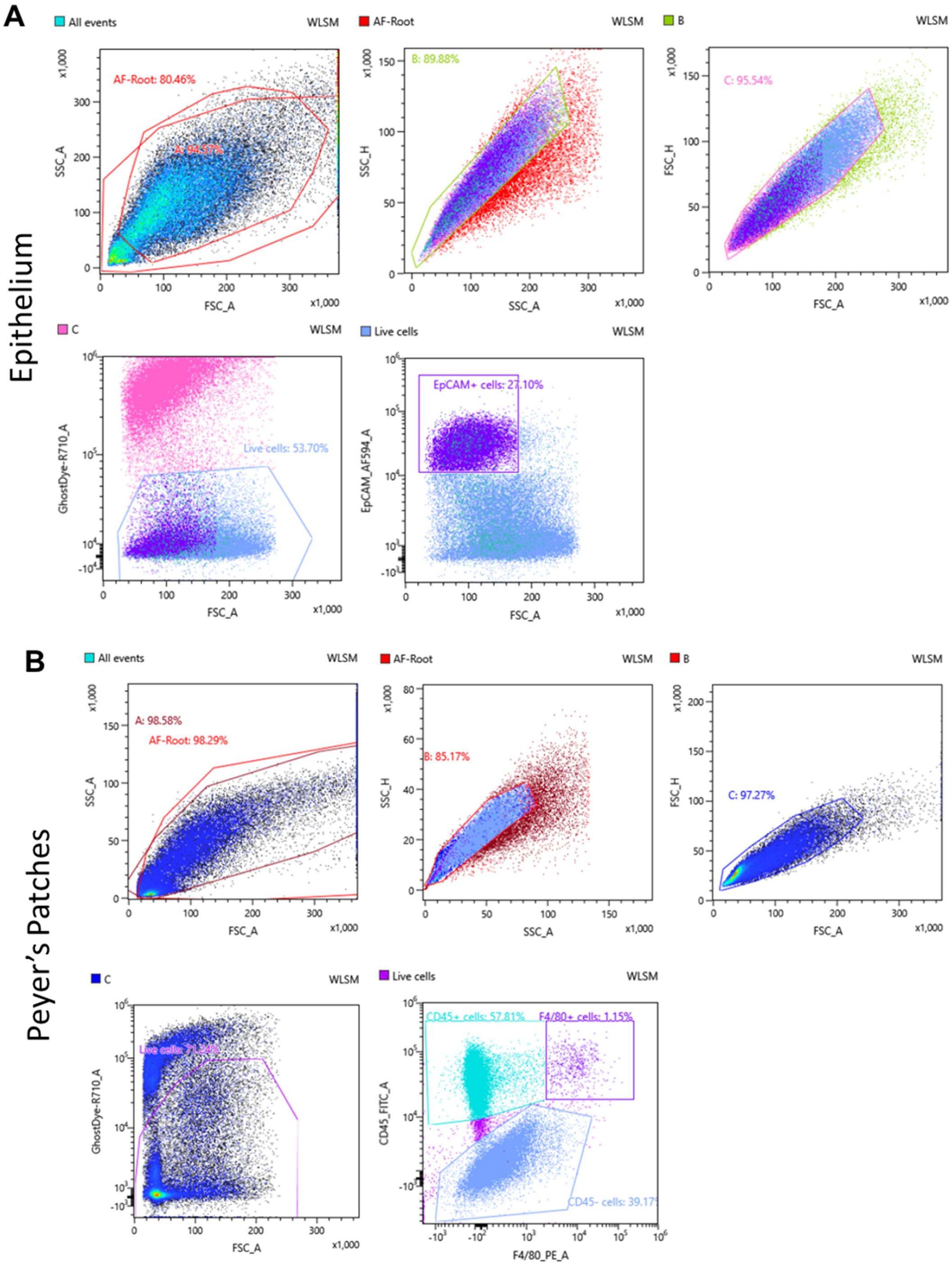

**C**

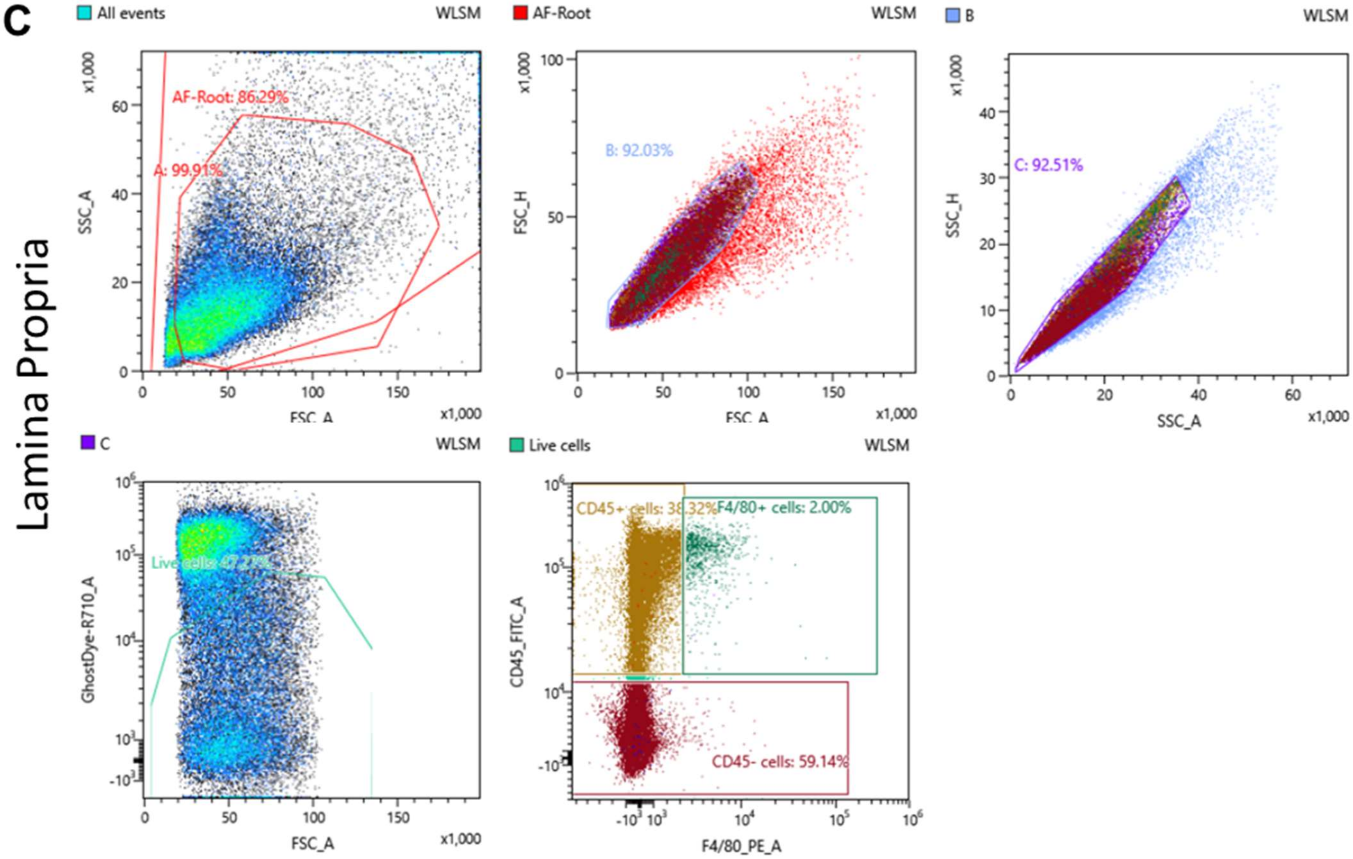
Lamina Propria
All events
WLSM
AF-Root: 86.29%
A: 99.91%
SSC_A
FSC_A
x1,000
AF-Root
B: 92.03%
FSC_H
B
C: 92.51%
SSC_H
C
Live cells: 47.27%
GhostDye-R710_A
Live cells
CD45+ cells: 38.32%
F4/80+ cells: 2.00%
CD45- cells: 59.14%
CD45_FITC_A
F4/80_PE_A
